# *Cryptococcus neoformans* employs both PP1 and PP2A phosphatases to silence the spindle checkpoint

**DOI:** 10.1101/2025.11.04.686612

**Authors:** Ardra Pamburayath Suresh, Haomiao Cheng, Thomas Davies, Ivan Clark, Christos Spanos, Kevin G. Hardwick

**Affiliations:** Institute of Cell Biology, School of Biological Sciences, University of Edinburgh, Edinburgh, EH9 3BF, UK

## Abstract

The spindle checkpoint preserves genomic integrity by delaying anaphase onset until all kinetochores achieve bi-polar microtubule attachment. While checkpoint activation is well-characterised across eukaryotes, the mechanisms governing its silencing remain less defined and are more variable in different systems. In this study, we reveal that the fungal pathogen *Cryptococcus neoformans* employs a dual phosphatase–mediated mechanism to silence the spindle checkpoint. We show that PP1 is recruited to kinetochores by Spc105^KNL1^, *via* conserved SILK and RVSF motifs, while PP2A-B56 is recruited to kinetochores through direct Bub1 binding *via* an LxxIxE motif. Disruption of either recruitment pathway leads to prolonged checkpoint signalling and mitotic defects, underscoring the critical roles of these phosphatases in checkpoint silencing and mitotic exit. These findings further establish *C. neoformans* as a powerful model for studying mitotic regulation in the basidiomycete lineage and identify phosphatase recruitment interfaces as potential targets for antifungal intervention.

## Introduction

Faithful chromosome segregation during mitosis is essential for maintaining genomic stability. To safeguard this process, eukaryotic cells employ the spindle checkpoint, a conserved surveillance mechanism that monitors attachment of spindle microtubules to kinetochores (McAinsh & Kops, 2023). If even a single kinetochore is unattached or incorrectly oriented, this checkpoint delays cell cycle progression at metaphase (Li & Nicklas, 1995; Rieder *et al*, 1995), by inhibiting Cdc20 (Hwang *et al*, 1998; Kim *et al*, 1998) and thereby the anaphase-promoting complex/cyclosome (APC/C). When active, this ubiquitin ligase polyubiquitinates securin and cyclin, labelling them for proteolytic destruction and thereby triggering sister chromatid separation and anaphase elongation (McAinsh & Kops, 2023).

Checkpoint activation is driven by a conserved network of proteins, including Mps1 kinase, Mad and Bub proteins, which together produce a diffusible “wait anaphase” signal by inhibiting Cdc20, the APC/C co-activator. This diffusible signal is often referred to as the mitotic checkpoint complex [MCC: typically Mad2-BubR1-Bub3-Cdc20 (Sudakin *et al*, 2001;Hardwick *et al*, 2000)]. The checkpoint delay provides time for Aurora B kinase, as part of the chromosomal passenger complex (CPC), to destabilise erroneous kinetochore-microtubule attachments, enabling error correction to take place (Ruchaud *et al*, 2007).

Once all chromosomes are correctly bi-oriented and under tension, checkpoint signalling must be rapidly silenced to permit timely anaphase onset (Lara-Gonzalez *et al*, 2021). This is achieved through removal of checkpoint proteins from kinetochores, disassembly of the mitotic checkpoint complex, and dephosphorylation of checkpoint components and targets by the mitotic phosphatases PP1 and PP2A (Espert *et al*, 2014; Foley & Kapoor, 2013). PP1 is recruited to KNL1 (Spc105 in *S.cerevisiae* and *C.neoformans*) *via* SILK and RVSF motifs to dephosphorylate MELT repeats and dismantle the checkpoint signalling platform (Espert *et al*., 2014; Rosenberg *et al*, 2011). PP2A-B56, recruited through BubR1, counteracts Aurora B activity and promotes stable microtubule attachment (Corno *et al*, 2023; Kruse *et al*, 2013). The precise coordination of checkpoint activation and silencing is critical for chromosome fidelity, and its dysregulation can result in chromosome mis-segregation, aneuploidy, and cell death.

Although the spindle checkpoint has been studied extensively in metazoans and ascomycete yeasts, little is known about its regulation in the basidiomycete lineage. *Cryptococcus neoformans* is an opportunistic fungal pathogen in this lineage and a World Health Organization critical-priority organism due to its high disease burden in immunocompromised patients, particularly those with HIV/AIDS [WHO, 2022 (Rajasingham *et al*, 2017)]. Treatment options are limited and antifungal resistance is rising (Gow *et al*, 2022), underscoring the need for new therapeutic targets. During infection, *Cryptococcus* displays striking morphological diversity, producing small “seed” cells (Brown & Ballou, 2024; Denham *et al*, 2022) and large, polyploid titan cells (Dambuza *et al*, 2018; Okagaki & Nielsen, 2012; Zaragoza & Nielsen, 2013). The polyploid titan cells continue to divide and produce viable haploid progeny. However, this reductive division can be error-prone, generating some aneuploid daughters (Gerstein *et al*, 2015). Such genetic diversity in the aneuploid offspring can help the pathogen adapt to stressful environments (Wertheimer *et al*, 2016). However, it is a major problem in the clinic as extra copies of chromosome 1 have been demonstrated to confer resistance to azole drugs (Wertheimer *et al*., 2016).

The molecular basis of checkpoint activation and silencing in *C. neoformans* remains largely unexplored. A genome-wide kinase-knockout screen implicated the Bub1 and Mps1 checkpoint components in fungal virulence (Lee *et al*, 2016). Recent work by our group confirmed conservation of core checkpoint proteins (Mps1, Mad1, Mad2) and showed that Mps1-dependent phosphorylation of Mad1 is essential for checkpoint function (Aktar *et al*, 2024). Analysis of CnBub1 and CnSgo1 has demonstrated conserved signalling mechanisms, but also some interesting differences from widely studied systems. For example, the Cn*BUB1* gene did not duplicate and so Bub1 carries out the functions of both Bub1 and Mad3/BubR1 (Leontiou *et al*, 2023). Both Bub1 kinase activity and Sgo1 are required to maintain prolonged checkpoint arrests, yet the bulk of CnSgo1 is found at spindle poles, rather than kinetochores (Leontiou *et al*., 2023; Polisetty *et al*, 2024).

In this study we investigate the roles of the conserved serine/threonine phosphatases PP1 and PP2A-B56 in checkpoint silencing in *C.neoformans*. Using CRISPR/Cas9-mediated genome-engineering, live-cell imaging and microfluidics, we show that PP1 is recruited to Spc105^KNL1^ *via* canonical SILK and RVSF motifs, while PP2A-B56 directly binds Bub1 through a conserved LxxIxE-like motif. We develop a synthetic spindle checkpoint assay, employing absiscic acid to bring together key signalling components (Mps1 and Spc105, or Mps1 and Bub1). Using this SynCheck assay, we demonstrate that disruption of the recruitment of either phosphatase prolongs checkpoint signalling and delays mitotic exit. Thus *Cryptococcus* employs a dual-phosphatase silencing mechanism, more like that of vertebrates than the ascomycete model yeasts *S.cerevisiae* and *S.pombe*.

## Results

### Mps1 kinase dependent phosphorylation of Spc105^KNL1^ is sufficient to activate the spindle checkpoint in *C. neoformans*

Several kinases and phosphatases play multiple mitotic roles, both in spindle checkpoint signalling and in the regulation of kinetochore-microtubule attachments (Moura *et al*, 2017;Corno *et al*, 2023). We wanted to dissect checkpoint activation and silencing mechanistically, in a way that would be independent of kinetochore-microtubule attachments. To do this, we adapted a synthetic checkpoint assay that we previously developed for fission yeast in which abscisic acid (ABA)-induced dimerisation brings the Mps1 kinase into close proximity with specific checkpoint substrates (Amin *et al*, 2018). In this assay the plant hormone abscisic acid (ABA) can be added to growth media to induce dimerisation of two proteins, tagged with the PYL and ABI signalling domains. The PYL fragment was fused to the N-terminal region of Spc105 (residues 1-600), which lacks the Mis12-binding site and so cannot localise to kinetochores (Petrovic *et al*, 2014). The ABI domain was fused to the C-terminal Mps1 kinase domain (residues 474–845 of Mps1), removing N-terminal targeting domains (Nijenhuis *et al*, 2013) and thereby preventing Mps1 kinetochore localisation (Fig. 1A). Both of these constructs were integrated at genomic safe haven loci and expressed from constitutive promoters for stable expression. Initially our strains carried an mCherry– Bub1 reporter to monitor checkpoint activation, when fluorescent Bub1 localises to clustered, mitotic kinetochores (Leontiou *et al*., 2023).

**Figure 1.**
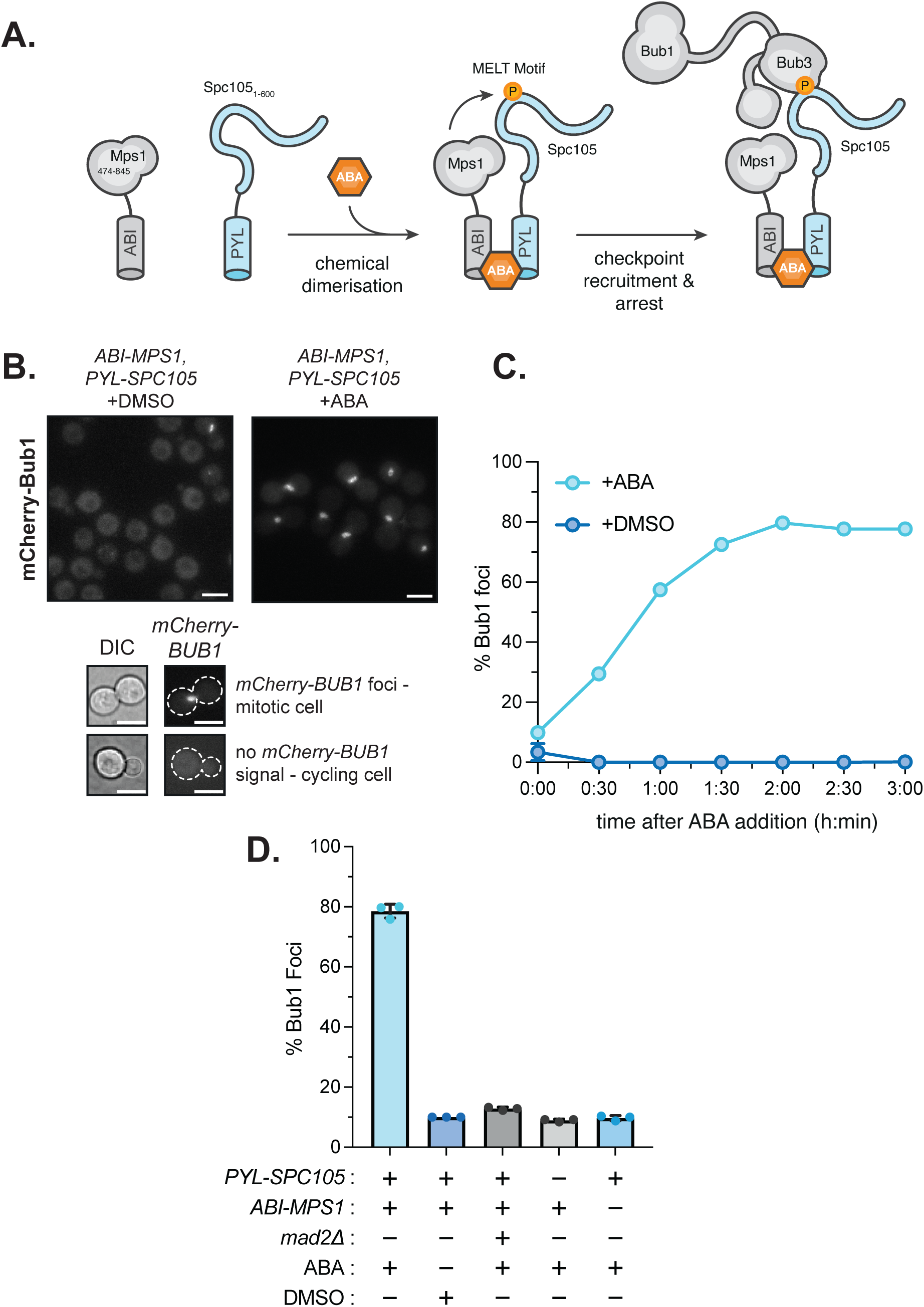
Establishment of the Mps1–Spc105 SynCheck system to induce metaphase arrest in *C. neoformans*. (A) Schematic representation of the synthetic checkpoint (SynCheck) assay. ABA-induced dimerisation between Mps1-ABI and Spc105-PYL enables phosphorylation of Spc105 MELT motifs, leading to recruitment of Bub1 and activation of the spindle checkpoint. (B) Representative fluorescence microscopy images showing mCherry-Bub1 recruitment to kinetochores. Scale bar, 5 µm. (C) Quantitation of the accumulation of cells in metaphase following ABA treatment in Spc105-PYL + Mps1-ABI cells. (D) Quantification of mCherry-Bub1 kinetochore localisation upon ABA treatment across different SynCheck configurations. Data represent mean ± s.d. from n = 3 independent experiments.

When ABA was added to cells expressing both Mps1-ABI and Spc105-PYL, Bub1–mCherry foci appeared within 30 minutes and a robust, large-budded, metaphase arrest (∼80% of cells) was observed after 2 hours (Fig. 1B&C). Importantly, this mitotic arrest required both components of SynCheck (Mps1-ABI and Spc105-PYL) and was abrogated in *mad2*Δ cells (Fig. 1D). We conclude that Mps1 kinase-dependent phosphorylation of the N-terminal fragment of Spc105 is sufficient to trigger checkpoint arrest in *C. neoformans*.

### Threonine 302 lies in a critical Spc105 MELT motif required for Mps1-dependent checkpoint activation

We next examined the role of MELT motifs in Spc105, which in other eukaryotes are phosphorylated by Mps1 to recruit Bub protein complexes to kinetochores. It is Bub3 that directly binds the phosphorylated MELT motif (Primorac *et al*, 2013) and this recruits both Bub3-Bub1 and Bub3-BubR1 complexes in human cells. The *C. neoformans* Spc105 sequence contains two canonical MELT motifs (MDIT^250^ and MDFT^302^) and one non-canonical MELT-like motif (MEET^356^) (Fig. 2B). We carried out *in vitro* kinase assays, employing recombinant Mps1 kinase (His-Sumo-CnMps1) and a fragment of Spc105 containing its first 600 residues (His-MBP-Spc105). Autoradiography demonstrates strong phosphorylation of the Spc105 fragment (Fig. 2A) and mass spec analysis of a non-radioactive kinase assay identified several phosphopeptides, one of which encompassed MDFT^302^ (Fig. 2B). 20% of MDFT peptides identified in this mass-spec analysis were phosphorylated, comparing favourably with phosphorylation levels of the N-terminal Spc105 sites we also identified (T26, T30, T45, T92, S96 and T104). To assess the relative contribution of the three MELT motifs, we generated a single alanine substitutions at T302A and a triple mutant (T250A, T302A, T356A) referred to here as T3A.

**Figure 2.**
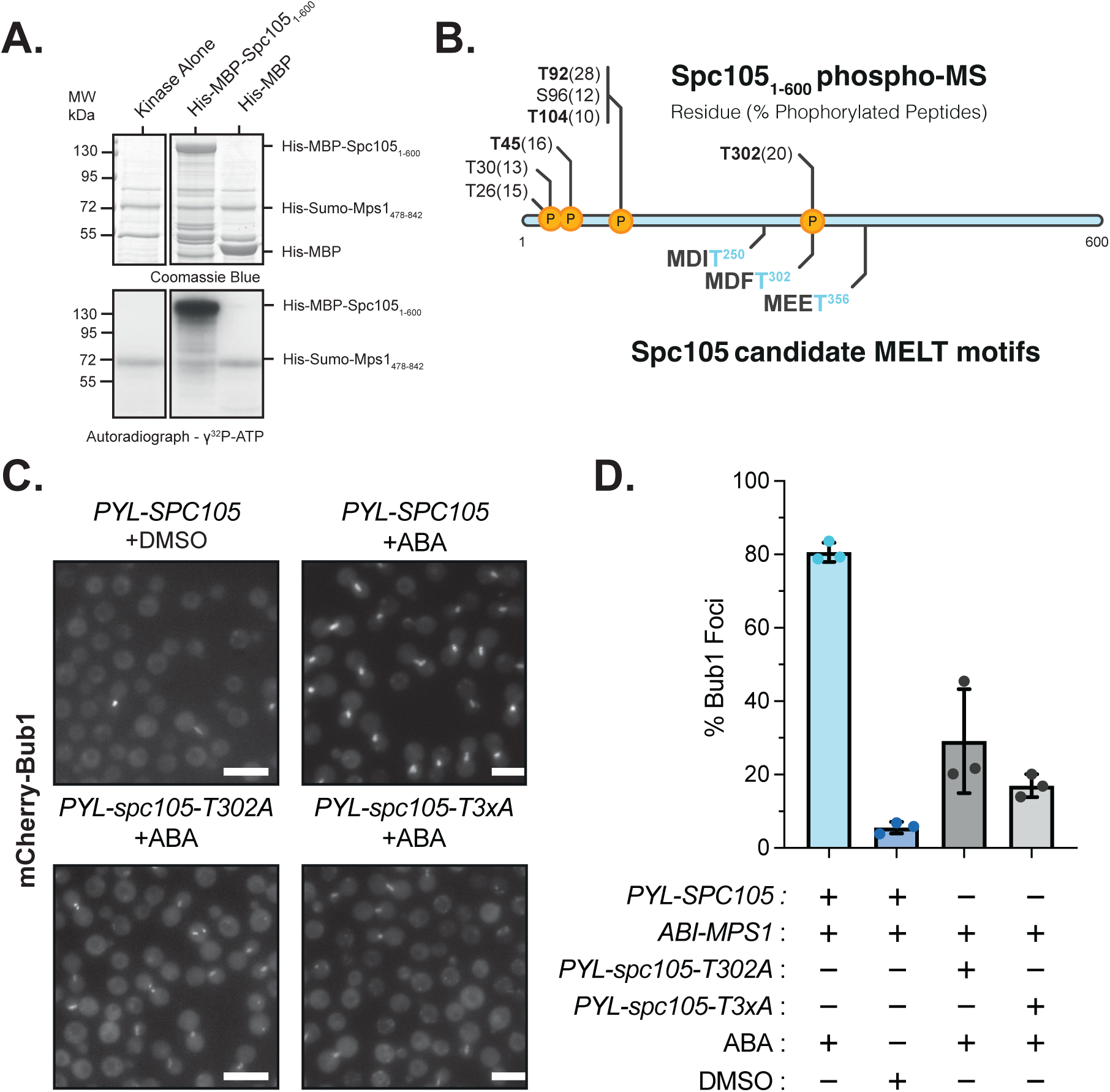
Functional analysis of Spc105 MELT motifs defines the minimal phosphorylation requirement for checkpoint activation. (A) *In vitro* kinase assays demonstrating Mps1 phosphorylation of Spc105 MELT motifs. Autoradiograph of ɣ^32^P-ATP (bottom) and corresponding Coomassie blue stained SDS-PAGE gel from ‘cold’ kinase assays (top). (B) Schematic of candidate MELT motif organisation (bottom) and corresponding phospho-mass spectrometry peptides (top) in Spc105. Phospho-peptides present in >10% of mass spectrometry hits are shown in approximate locations within the Spc105 amino acid sequence. (C) Representative fluorescence microscopy images showing mCherry-Bub1 recruitment to kinetochores in cells expressing Spc105 variants following ABA treatment. Scale bar 10 µm. (D) Quantitation of metaphase arrest efficiency for the indicated constructs after 2 h of ABA exposure. Data represent mean ± s.d. from *n* = 3 independent experiments.

Both the single (T302A) and the triple mutant significantly impaired the SynCheck arrest (Fig.2C-D). In wild-type SynCheck strains ∼80% of cells displaying mCherry-Bub1 foci after two hours of ABA treatment for two hours, yet only ∼30% of cells arrested in the *spc105*-*T302A* mutant and ∼17% in the *spc105*-*3A* mutant (Fig.2D). It was noticeable that the mutants both displayed weaker and more fragmented Bub1 foci. These data point to a dominant role for the T302 MELT motif, with T250 and/or T356 making a smaller but measurable contribution. These results confirm that Mps1-dependent MELT phosphorylation of Spc105 is sufficient to trigger checkpoint arrest.

### Spc105 can be by-passed via direct phosphorylation of Bub1 in SynCheck-ABA

We next asked whether checkpoint activation could be achieved by directly targeting phosphorylation of Bub1 by Mps1 kinase, bypassing the need for Spc105. *Cryptococcus* Bub1 is also an active checkpoint kinase (Leontiou *et al*., 2023), so to simplify our Mps1-Bub1 SynCheck assay, we used a Bub1-PYL construct containing residues 1–899 of Bub1 but lacking the C-terminal kinase domain (Fig. 3A). Previous work in ascomycete yeasts and metazoans has shown that Mps1 phosphorylates Bub1 within a conserved domain known as CD1, enabling it to directly bind and recruit Mad1-Mad2 complexes (Brady & Hardwick, 2000; Fischer *et al*, 2021; Heinrich *et al*, 2014; Klebig *et al*, 2009). Our previous work (Leontiou *et al*., 2023) has established that *C. neoformans* Bub1 is a MadBub (Vleugel *et al*, 2012) and functions as a central signalling hub, combining the upstream signalling scaffold role of Bub1 with the downstream APC/C-inhibitory role typically performed by BubR1/Mad3. Thus CnBub1 integrates upstream kinase signalling (by CDK and Mps1) with direct inhibition of anaphase onset. We have previously shown that mutation of CnBub1-CD1, the conserved domain of Bub1 that gets phosphorylated by both CDK1 and Mps1 to enable Mad1 binding, completely abolished the *Cryptococcus* checkpoint response (Leontiou *et al*., 2023), as did mutation of the first KEN box in CnBub1.

**Figure 3.**
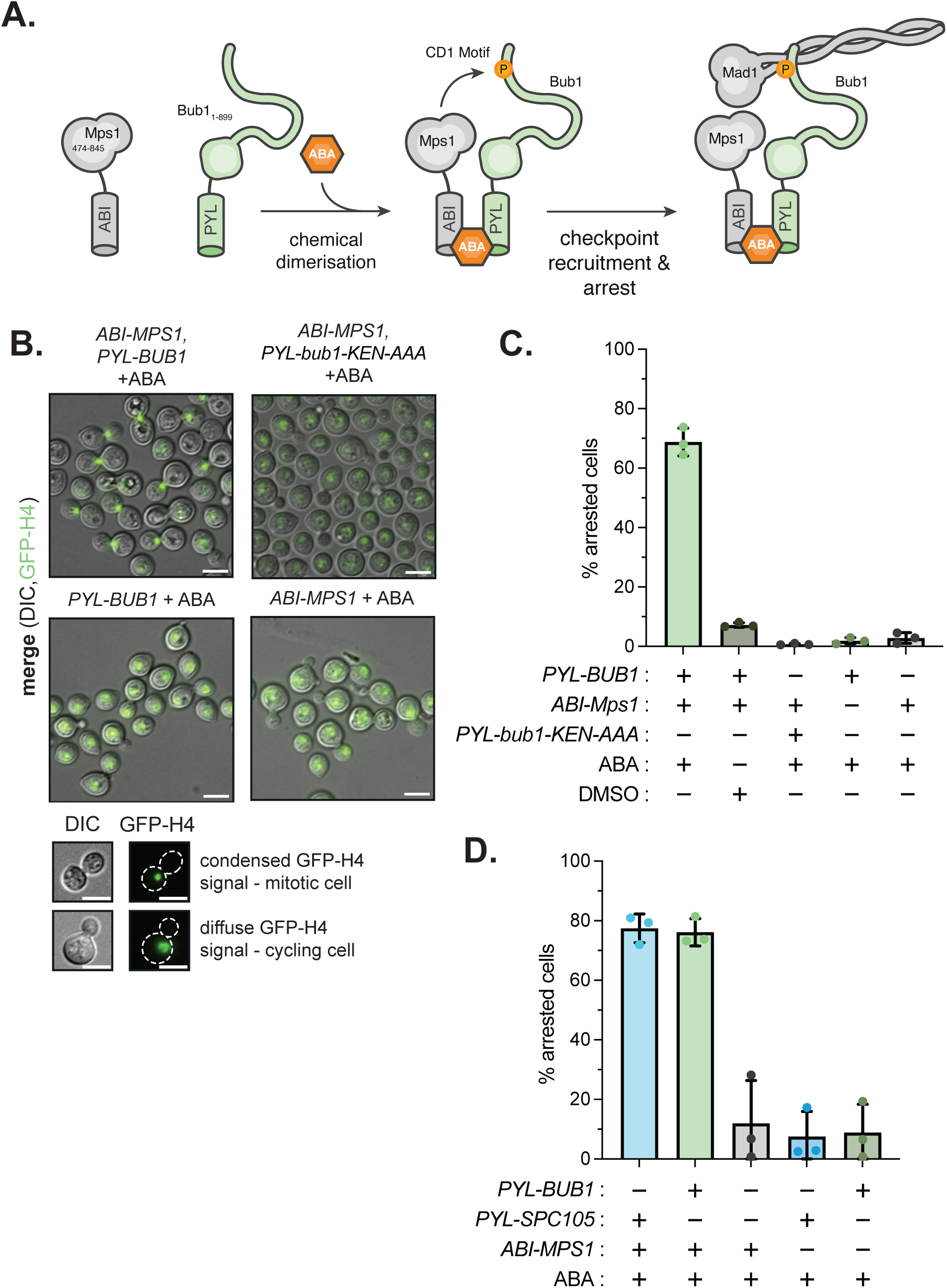
Spc105 can be by-passed in the Mps1–Bub1 SynCheck assay. (A) Schematic of the Mps1–Bub1 SynCheck system. ABA-induced dimerisation between Mps1-ABI and Bub1-PYL activates the checkpoint, with metaphase arrest monitored using a histone H4 reporter to visualise nuclear morphology. (B) Representative microscopy images showing metaphase arrest - scored *via* condensed GFP-H4 signal - in cells carrying the Bub1–Mps1 SynCheck system. Scale bar, 5 µm. (C) Quantitation of metaphase arrest efficiency in wild-type (H99) Bub1-Mps1 versus *bub1-ken1*-Mps1 and control strains. Data represent mean ± s.d. from *n* = 3 independent experiments. (d) Comparison of arrest efficiency between the Spc105–Mps1 and Bub1–Mps1 SynCheck systems after 2 h ABA treatment. Data represent mean ± s.d. from *n* = 3 independent experiments.

Here we used our SynCheck assay to test whether direct phosphorylation of Bub1 by Mps1 is sufficient to trigger checkpoint arrest, bypassing the Spc105 kinetochore-based signalling platform. As we are dissecting Bub1 functions, we replaced our Cherry-Bub1 reporter with GFP-Histone H4 for the remaining SynCheck experiments. After two hours of ABA treatment, strains expressing both Mps1-ABI and Bub1-PYL produced robust metaphase arrest, with 70-80% of cells being large-budded with condensed GFP-H4 signals (Fig. 3B&C). This arrest was ABA-dependent, and neither Bub1-PYL nor Mps1-ABI were sufficient on their own to generate an arrest (Fig. 3B&C). Importantly, mutation of the N-terminal KEN box in the Bub1-PYL construct completely abolished the SynCheck arrest. This KEN box binds Cdc20 when HsBubR1 forms the MCC complex (Alfieri *et al*, 2016), and we have demonstrated that it is critical for arrest in CnBub1 (Leontiou *et al*., 2023). Thus, this SynCheck assay appears to be accurately reflecting normal signalling mechanisms. We wanted to test how efficient this Mps1-Bub1 arrest was, when compared to the Mps1-Spc105-PYL assay in Figs. 1&2. Fig. 3D shows that the Bub1-PYL arrests cells with a very similar efficiency to that of Spc105-PYL. Our interpretation is that *Cryptococcus* Bub1 is itself sufficient to form a highly efficient checkpoint signalling scaffold, once it is phosphorylated by Mps1 kinase. Whilst Spc105 MELT motifs normally form the kinetochore-receptor for the Bub1 checkpoint signalling platform, they are not needed in our diffusible SynCheck assay.

### Dual PP1 and PP2A-B56 phosphatase pathways drive spindle checkpoint silencing in *C. neoformans*

Having established two orthogonal routes to trigger a strong and sustained spindle checkpoint arrest, we next asked whether *Cryptococcus* cells could recover from this metaphase block, and if so what silencing mechanisms were employed. Here we focus on the contributions of phosphatases. In metazoans, spindle checkpoint silencing is mediated by the coordinated action of PP1 and PP2A-B56 (Espert *et al*., 2014; Smith *et al*, 2019). In contrast, studies in the model ascomycete yeasts *S.cerevisiae* and *S. pombe* have shown that PP1 is the major player in checkpoint silencing (Meadows *et al*, 2011; Pinsky *et al*, 2009; Rosenberg *et al*., 2011; Vanoosthuyse & Hardwick, 2009).

We extended our SynCheck assay with ABA washout to terminate checkpoint activation (Fig. 4A). This experimental design allowed us to measure the kinetics of checkpoint inactivation without the confounding effects of kinetochore-microtubule attachment defects that are found in mitotic blocks such as nocodazole arrest. PP1 is predicted to bind directly to Spc105-PYL, *via* conserved SILK and RVSF motifs (Fig. 4B). PP2A is predicted to bind directly to Bub1-PYL *via* an LxxIxE-like motif, MTP**I**T**E** (Fig. 4B). ABA was washed out after 2 hours of arrest, and both the Mps1-ABI, Spc105-PYL and the Mps1-ABI, Bub1-PYL wild-type strains showed rapid recovery: the proportion of metaphase-arrested cells dropped from ∼80% at the point of washout to ∼30% within 30 mins, and <10% by 60 mins (Fig. 4C&D). This is interesting as it demonstrates that both phosphatases are involved in checkpoint silencing in *Cryptococcus*, resembling metazoans rather than model ascomycetes. As the Mps1-ABI, Bub1-PYL Syncheck bypasses Spc105, we conclude that PP2A is sufficient to inactivate/silence this Bub1-PYL signalling scaffold.

**Figure 4.**
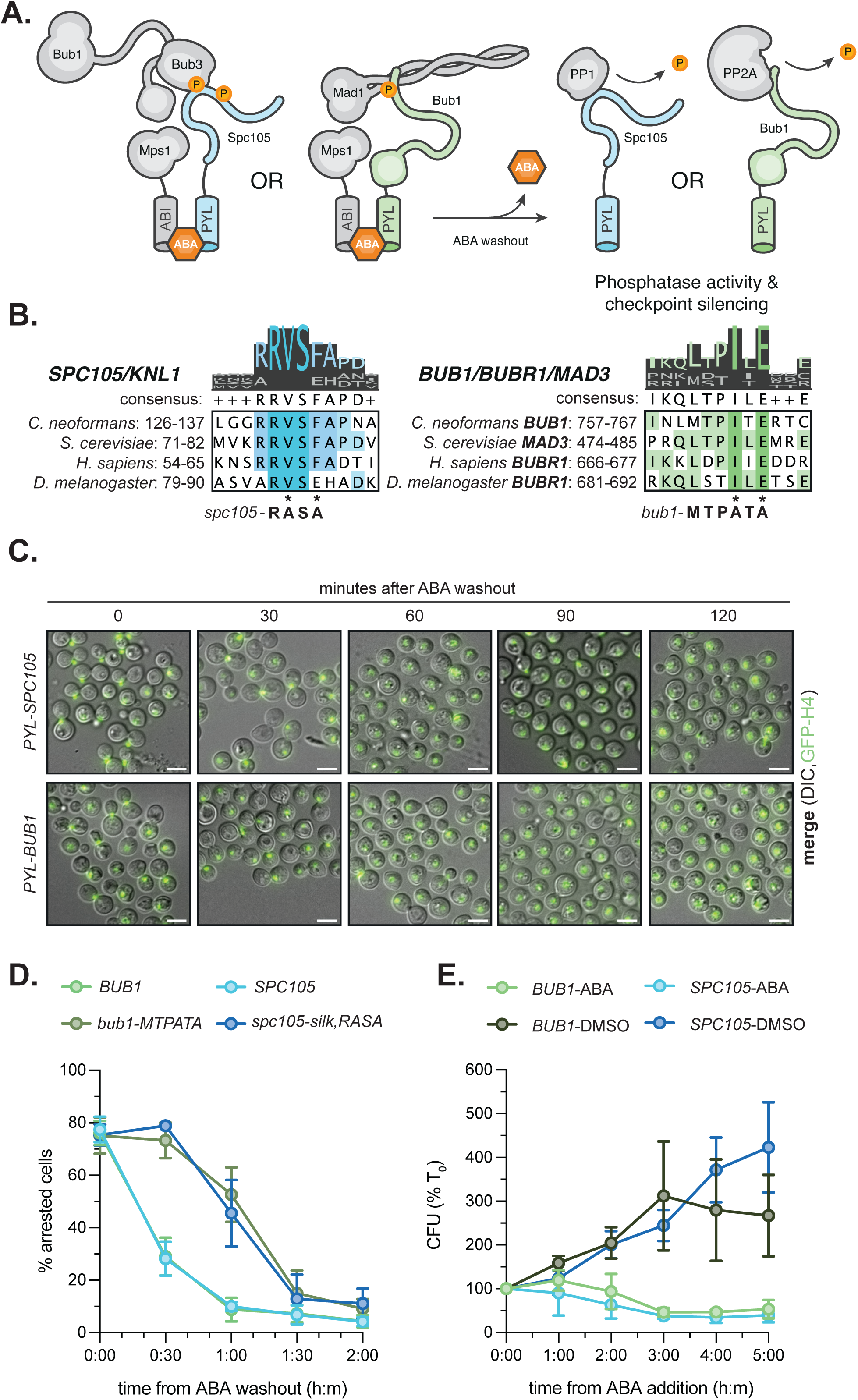
Both PP1- and PP2A-dependent checkpoint silencing pathways mediate recovery from *Cryptococcus* SynCheck arrests. (A) Schematic of the SynCheck washout assay used to monitor checkpoint silencing in the Spc105–Mps1 and Bub1–Mps1 systems. Cells are arrested in metaphase by ABA-induced dimerisation and then released by ABA removal. Phosphatase activity then leads to checkpoint silencing. (B) Multiple sequence alignment of conserved RVSF motifs in Spc105 (left) and LxxIxE motifs in Bub1 (right), in *C. neoformans*, *S. cerevisiae*, *H. sapiens*, and *D. melanogaster* homologues, generated in Jalview. (C) Representative microscopy images of wild-type Spc105–Mps1 and Bub1–Mps1 SynCheck strains at 0, 30, 60, 90, and 120 min following ABA washout, showing progressive recovery from arrest. Scale bar, 5 µm. (D) Quantification of the proportion of cells remaining in metaphase following ABA washout across different genotypes. Data represent mean ± s.d., n = 3 independent experiments. (E) Cell viability after 0-5h of ABA incubation. Wild-type Spc105-PYL–Mps1-ABI and Bub1-PYL– Mps1-ABI strains were arrested with ABA or treated with DMSO for 0-5 h, washed remove ABA, and plated at 0, 1, 2, 3, 4, 5 h after adding ABA or DMSO. Data represent mean ± s.d., n = 3 independent experiments.

In many systems, prolonged mitotic arrests have been demonstrated to lead to a decrease in viability (Amin *et al*., 2018; Gascoigne & Taylor, 2008). We were hoping that this would not be the case in SynCheck. If so, we might be able to use SynCheck to synchronise *Cryptococcus* cultures in mitosis, something that is currently very difficult to achieve. Both SynCheck strains were arrested for different times in ABA, carefully washed and then plated for colony-forming viability assays (Fig. 4E). This revealed that cells arrested for prolonged periods (>2 hours) exhibited a progressive decline in plating efficiency compared to untreated controls. This closely matches the viability loss observed after treatment of *Cryptococcus* with the anti-microtubule drug nocodazole (data not shown). It is also consistent with previous SynCheck studies in *S. pombe*, where extended synthetic arrest caused a time-dependent loss of viability (Amin *et al*., 2018; Yuan *et al*, 2017). More work is needed to see if a 90 min ABA block/release can be used to reproducibly synchronise viable *Cryptococcus* cultures. Future studies will investigate why *Cryptococcus* cells die after prolonged ABA arrest.

To dissect the contribution of each phosphatase, we generated strains with targeted mutations in the known phosphatase-recruitment motifs. In Spc105–PYL we mutated the conserved RVSF motif to RASA and SILK to AAAA, preventing PP1 docking (Rosenberg *et al*., 2011). In Bub1–PYL we mutated the conserved hydrophobic isoleucine (I) residue and the glutamic acid (E) in the LxxIxE-like motif in CnBub1, producing the *bub1-*MTPATA mutant to prevent PP2A binding (Hertz *et al*, 2016; Suijkerbuijk *et al*, 2012). In both cases, mutation of the phosphatase docking site caused a pronounced delay in silencing with ∼50% of cells remaining arrested 60 mins after ABA removal (Fig. 4D). We conclude that both PP1 (bound to Spc105) and PP2A-B56 (bound to Bub1) are critical for timely checkpoint silencing in *C.neoformans*.

### Bub1 recruits PP2A-B56 to kinetochores in *C. neoformans*

This involvement of PP2A-B56, recruited *via* CnBub1, is particularly novel and interesting. We next wanted to analyse its role in normal mitotic regulation, independent of SynCheck. To do this we characterised the Bub1-PP2A interaction *in vivo* using both co-immunoprecipitation and co-localisation studies. To test whether this motif mediates PP2A-B56 binding, we complemented a *bub1*1 strain with GFP-Bub1-MTPATA expressed from an ectopic, safe haven locus in a strain also expressing mCherry-tagged PP2A. Co-immunoprecipitation experiments using GFP-Trap beads revealed that in wild-type cells, Bub1 strongly associated with PP2A-B56 in cell lysates, both in the absence and the presence of nocodazole (Fig. 5A). This suggests the Bub1-PP2A complex is constitutive, rather than being cell cycle regulated. Mutation of the MTPITE motif completely abolished this interaction. Future experiments will test whether the Bub1-PP2A interaction is regulated by Polo kinase, as found in vertebrate cells (Corno *et al*., 2023).

**Figure 5.**
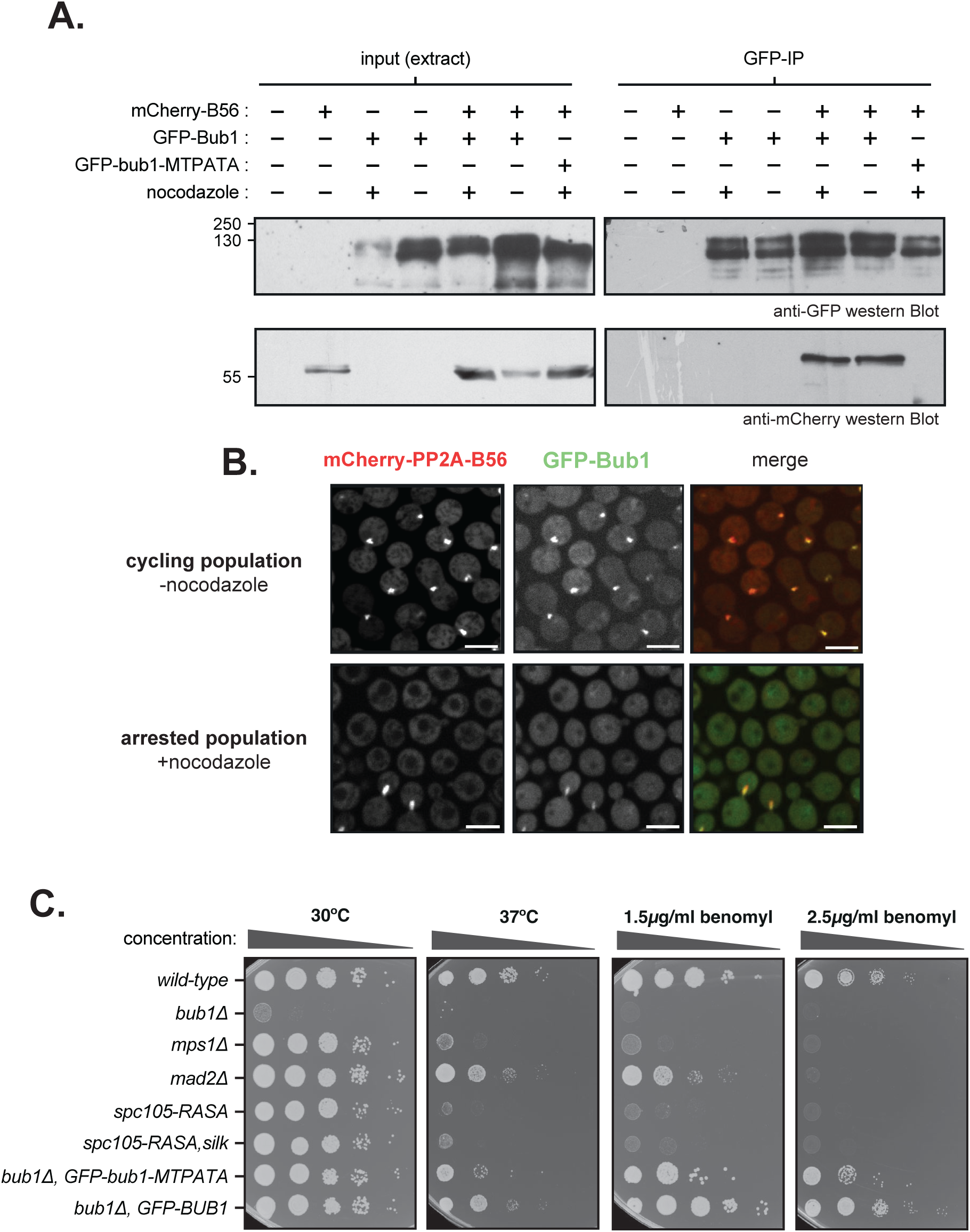
PP2A is recruited to CnBub1 via a conserved LxxIxE motif. (A) Representative co-immunoprecipitation of GFP-Bub1 with mCherry-PP2A-B56γ. Native protein lysates were prepared from the indicated strains, incubated with GFP-TRAP beads, washed, and bound proteins immunoblotted with anti-GFP and anti-mCherry antibodies. Input of whole cell extract and IP are shown. The experiment was repeated three times with similar results. (B) Representative microscopy images showing colocalization of PP2A-B56γ with Bub1 at kinetochores during checkpoint arrest. Scale bar, 5 µm. (C) Representative serial dilution growth assays of strains carrying endogenous mutations in the Spc105 RVSF motif (PP1-binding) and the Bub1 LxxIxE motif (PP2A-B56γ-binding). Cells were spotted onto YPD with or without benomyl and incubated at either 30 °C or elevated temperature (37 °C) to assess sensitivity to spindle and temperature stress.

We next assessed PP2A localisation *in vivo*. Fluorescence microscopy showed that mCherry-B56 accumulated at kinetochores during checkpoint arrest, colocalising with GFP-Bub1 foci (Fig. 5B). In the *bub1-*MPTATA mutant, GFP-Bub1 foci were retained but mCherry-B56 failed to co-localise, remaining diffuse in the nucleoplasm and cytoplasm (not shown). This loss of kinetochore enrichment mirrored the lack of co-immunoprecipitation and suggests that Bub1 directly recruits PP2A to mitotic kinetochores through the MTPITE motif.

Taken together, these results establish that *C. neoformans* Bub1 serves as a direct scaffold for PP2A-B56 recruitment during checkpoint arrest, using a conserved SLiM that is functionally equivalent to that described in metazoan BubR1 and Bub1 (Kruse *et al*., 2013); Xu et al., 2013). This dual role for Bub1, promoting checkpoint activation whilst simultaneously engaging PP2A-B56 for silencing, highlights CnBub1 as a central node in mitotic regulation. The finding that basidiomycete Bub1 retains this dual functionality suggests that the double phosphatase system is an ancient feature of eukaryotic spindle checkpoint control, rather than a metazoan innovation.

### Mutations in PP1 and PP2A recruitment motifs impair checkpoint silencing and growth/viability

Our SynCheck and biochemical experiments established that both PP1 and PP2A-B56 are required for efficient spindle checkpoint silencing in *C. neoformans*. To test whether these requirements hold true in their native context, we generated strains carrying endogenous mutations in the phosphatase-docking motifs of Spc105 and Bub1. Specifically, we replaced the canonical SILK and RVSF motifs of Spc105, which mediates PP1 recruitment, with alanines (SILK-AAAA, RVSF-RASA) and introduced point mutations into the conserved LxxIxE-like motif of Bub1 (MTPATA).

Mutation of individual phosphatase-docking sites produced strains that were viable under standard growth conditions (room temperature on YPDA plates), indicating that basal mitotic progression does not strictly depend on either individual pathway. However, when cells were challenged with spindle stress (benomyl treatment), their deficiencies became evident. *spc105-RASA, spc105-silk,RASA* and *bub1-MTPATA* mutants all displayed strong growth defects on benomyl-containing medium and at high temperature (Fig. 5C). This stress sensitivity highlights the importance of efficient checkpoint silencing in enabling survival when kinetochore-microtubule attachments are perturbed.

Finally, and to quantitatively assess checkpoint dynamics at the single-cell level, we employed microfluidics-based, live-cell imaging to monitor responses to nocodazole over time. Strains were pre-grown in minimal (SC) medium and loaded into the microfluidic device, where individual cells were captured and imaged prior to, during and after drug treatment. Nocodazole (2.5 μg/ml) was added after 5 hrs, and imaging was continued for an additional 3 hrs at 2 min intervals. Under these conditions, all strains except the checkpoint-defective *mad2*1 strain, arrested in metaphase with bright GFP-Bub1 foci at unattached kinetochores. Following drug washout, cells were imaged for a further 12 hrs, allowing us to track their recovery and subsequent divisions. Movies were analysed to quantify the proportion of cells that arrested upon nocodazole treatment and the fraction that successfully re-budded and resumed division after release. The wild-type cells typically arrested in metaphase upon nocodozole addition and exited the arrest within 3 hours of washout. In contrast, both phosphatase-docking mutants exhibited markedly delayed recovery, remaining in metaphase much longer after removal of nocodazole (Fig. 6A). Quantitative analysis of GFP-Bub1 signals confirmed that loss of either PP1 recruitment to Spc105 or PP2A-B56 recruitment to Bub1 substantially delayed checkpoint silencing by around 30 mins, as assessed by delayed loss of GFP-Bub1 foci (Fig. 6B).

**Figure 6.**
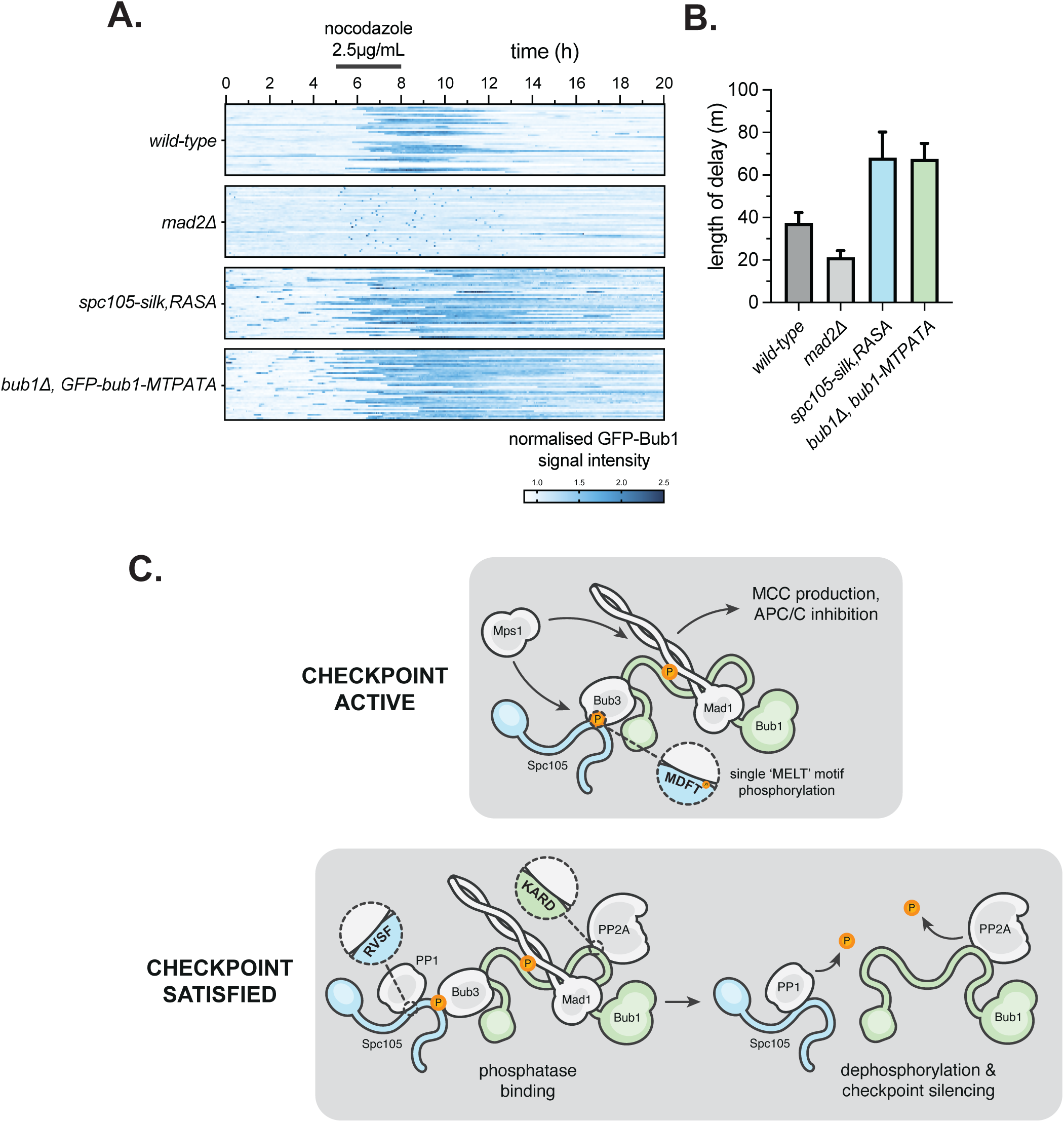
Microfluidics reveals delayed checkpoint silencing when kinetochore recruitment of either PP1 or PP2A is impaired. (A) Representative kymographs (30 per strain) showing GFP–Bub1 kinetochore localisation in wild type, *bub1-cd1*, and phosphatase recruitment mutants (*spc105-RASA* and *bub1-MTPATA*) are shown. Bright GFP–Bub1 signals (>1.22 normalised GFP-Bub1 intensity) mark mitosis, reflecting Bub1 enrichment at kinetochores. Nocodazole was present from 5 to 8 hrs (indicated by grey line above plots) but then washed out to analyse checkpoint silencing. (B) Quantitation of the average time spent with bright GFP-Bub1 at kinetochores, after nocodazole wash-out. (C) Schematic model illustrating phosphatase-dependent silencing pathways. PP1 is recruited via the Spc105 RVSF motif and PP2A via the Bub1 MxxIxE motif. Both phosphatases contribute to checkpoint inactivation following kinetochore attachment: disruption of either recruitment pathway results in delayed silencing and prolonged metaphase arrest.

Taken together, these results demonstrate that both PP1 and PP2A-B56 pathways contribute to spindle checkpoint silencing *in vivo*. While cells can tolerate disruption of either pathway under normal conditions, both become essential for viability under spindle stress, when efficient recovery from checkpoint arrest is critical. Silencing of the spindle checkpoint in *C. neoformans* therefore requires the coordinated action of two phosphatases, resembling the situation in metazoans and diverging from the simplified PP1-only silencing mechanism of ascomycete yeasts (Fig. 6C). We predict that the *spc105-RASA, bub1-MTPATA* mutant will be dead (due to a lethal metaphase block): indeed numerous attempts to build such a strain have thus far proved unsuccessful.

## Discussion

Several distinct mechanisms have evolved to efficiently silence the checkpoint once all chromosomes have achieved bi-orientation (Lara-Gonzalez *et al*., 2021). This study establishes that the basidiomycete fungal pathogen *Cryptococcus neoformans* employs a dual phosphatase–mediated silencing system involving both PP1 and PP2A-B56. This more closely resembles metazoan mechanisms (Smith *et al*., 2019) than those described in ascomycete yeasts, which rely very heavily on PP1 (London *et al*, 2012; Meadows *et al*., 2011; Vanoosthuyse & Hardwick, 2009). Using synthetic checkpoint reconstitution, we have demonstrated that PP1 is recruited to the kinetochore protein Spc105^KNL1^ *via* canonical SILK/RVSF motifs, whilst PP2A-B56 is targeted to kinetochores *via* an LxxIxE-like motif on Bub1. Mutation of either docking site delays checkpoint silencing, and we predict that disruption of both recruitment mechanisms will be synthetically lethal.

The relevant mitotic substrates of CnPP1 and CnPP2A remain to be determined. In metazoans, PP1 primarily reverses Mps1-dependent phosphorylation of MELT repeats to dismantle the Bub1–Bub3 scaffold. PP2A-B56 aids in this process and also counteracts Aurora B phosphorylation to stabilise microtubule attachments (Corno *et al*., 2023; Espert *et al*., 2014; Moura *et al*, 2017).

From a mechanistic standpoint, our work expands understanding of spindle checkpoint silencing interfaces in fungi. *In vivo* co-immunoprecipitation and colocalisation experiments confirm that Bub1 acts as the primary kinetochore receptor for PP2A-B56 during checkpoint arrest. Interestingly this biochemical Bub1-PP2A interaction appears to be constitutive, so once Bub1 is targeted to mitotic kinetochores it will bring PP2A with it (Fig. 5). Both the PP2A and the PP1 binding sites are regulated by phosphorylation in vertebrate cells, by Plk1 and Aurora B respectively (Corno *et al*., 2023; Liu *et al*, 2010). It will be fascinating to determine whether such phospho-regulation of phosphatase recruitment is found in *Cryptococcus*.

The functional consequences of disrupting these phosphatase recruitment motifs in *C.neoformans* are profound. The *spc105-RASA* mutant and the *bub1-MTPATA* mutant both compromise recovery from checkpoint arrest, as seen in our live-cell microfluidics assays (Fig. 6A). This impairs their growth under spindle stress, such as on plates containing benomyl (Fig. 5C). These mutants are also very sensitive to elevated temperatures, as are *bub1*1 and *mps1*1 strains (Fig. 5C). The reasons for these high levels of sensitivity are as yet unclear, but likely relate to additional phosphatase functions at kinetochores, such as the stabilisation of kinetochore-microtubule attachments. Whilst these are not relevant in SynCheck assays (Figs. 1-4) they will be critical for the co-ordination of error-correction processes and spindle checkpoint silencing.

A key next step will be to construct a double mutant in which both PP1 and PP2A-B56 recruitment are simultaneously disrupted. We expect to see synthetic lethality when *spc105-RASA* and *bub1-MTPATA* mutants are combined. Indeed, when we attempted *via* CRISPR to mutate the PP1-binding motifs in Spc105 in the *bub1-MTPATA* background, we were unable to recover viable strains. Although technically challenging, establishing such a double mutant remains a key experiment, as it would provide definitive genetic evidence that the two phosphatases act in essential, overlapping pathways to terminate checkpoint signalling, even in a ‘normal’ unperturbed mitosis.

From an evolutionary perspective, the presence of both PP1 and PP2A-B56 silencing pathways in *C. neoformans* raises intriguing questions about how the checkpoint has adapted to different cellular architectures and ecological niches. Basidiomycetes like *C. neoformans* experience unique challenges, including dramatic morphological transitions such as titanisation and polyploidy. The reductive divisions, when polyploid titans produce haploid daughter cells are particularly challenging. Due to the number of chromosomes and spindle geometry it is likely that inappropriate kinetochore attachments (likely merotelics) will initially be made, and checkpoint and error-correcting kinases and phosphatases will become even more important for accurate chromosome segregation. This has been demonstrated in ascomycete model yeasts with increased ploidy (Storchova *et al*, 2011; Storchova *et al*, 2006). Such complexities may necessitate more layered silencing systems to maintain viability during prolonged mitoses and/or unconventional division cycles.

Finally, these insights may bear direct relevance to pathogenicity and the identification of future drug targets. Titan cells, a hallmark of *C. neoformans* infection, are polyploid and undergo error-prone divisions that can generate aneuploid progeny, fuelling antifungal drug resistance and immune evasion (Dambuza *et al*., 2018; Gerstein *et al*., 2015). We have observed that PP2A-B56 recruitment-defective mutants (*bub1-MTPATA*) show reduced titan cell viability (unpublished data), suggesting that this phosphatase pathway is particularly important for the stability of this morphotype. Impaired silencing in titan cells may exacerbate chromosome segregation errors, leading to non-viable or genetically unstable progeny. We are currently investigating whether this defect accelerates the emergence of aneuploid, drug-resistant subpopulations.

## Acknowledgements

We would like to thank all members of the Hardwick and JP labs for their support, discussions and suggestions on this manuscript. We thank Dave Kelly, Toni McHugh, Dhanya Cheerambathur and Peter Swain for their help with imaging, scripts and their analysis. We thank James Fraser and Paige Erpf for *Cryptococcus* safe haven and Blaster constructs; and the Hiten Madhani lab for assistance with CRISPR engineering.

This work was supported by grants from the Leverhulme Trust [RPG-2018-379, KH]; the Darwin Trust of Edinburgh [AS and HC] and the Wellcome Trust [Discovery Award 309153, KH & TD]. This work was also supported by funding for the Wellcome Discovery Research Platform for Hidden Cell Biology [226791] and we gratefully acknowledge support from the Light Microscopy and Proteomics cores. Our funders had no role in study design, data collection and analysis, decision to publish, or preparation of this manuscript.

## Methods

### Yeast media

All *Cryptococcus* strains were derived from the H99L genetic background and preserved in YPDA medium (2% bactopeptone, 1% yeast extract, 2% glucose, 2% agar, 1 mM adenine) supplemented with 30% glycerol at −70 °C. Strains were streaked onto YPDA plates and grown for two days before use. For microfluidic experiments, cells were cultured in synthetic complete medium (SC: 0.67% yeast nitrogen base, amino acid mix, 2% glucose). Knockout transformants generated with the blaster cassette were selected on YNB plates (0.45% yeast nitrogen base without amino acids and ammonium sulphate, 2% glucose, 10 mM ammonium sulphate, 2% agar, 1 mM adenine) supplemented with 5 mM acetamide. To excise the blaster cassette, strains were re-streaked on YNB plates containing 2% glucose, 10 mM ammonium sulphate, and 10 mM fluoroacetamide, as previously described (Erpf *et al*, 2019). For drug selection, cells were grown on YPDA plates supplemented with 300 µg/ml hygromycin B (Invitrogen), 100 µg/ml Geneticin (G418 sulphate, Gibco), or 100 µg/ml nourseothricin (Jena Bioscience).

#### Gibson assembly and sequencing

Primers were purchased from Sigma/Merck or IDT. Gibson and NEBuilder assembly reactions were carried out according to the manufacturer’s instructions. DNA sequencing was performed using BigDye chemistry.

### Strains used in this study

***Cryptococcus* strains** were all derived from *Cryptococcus neoformans* var. *grubii* H99L unless otherwise stated.

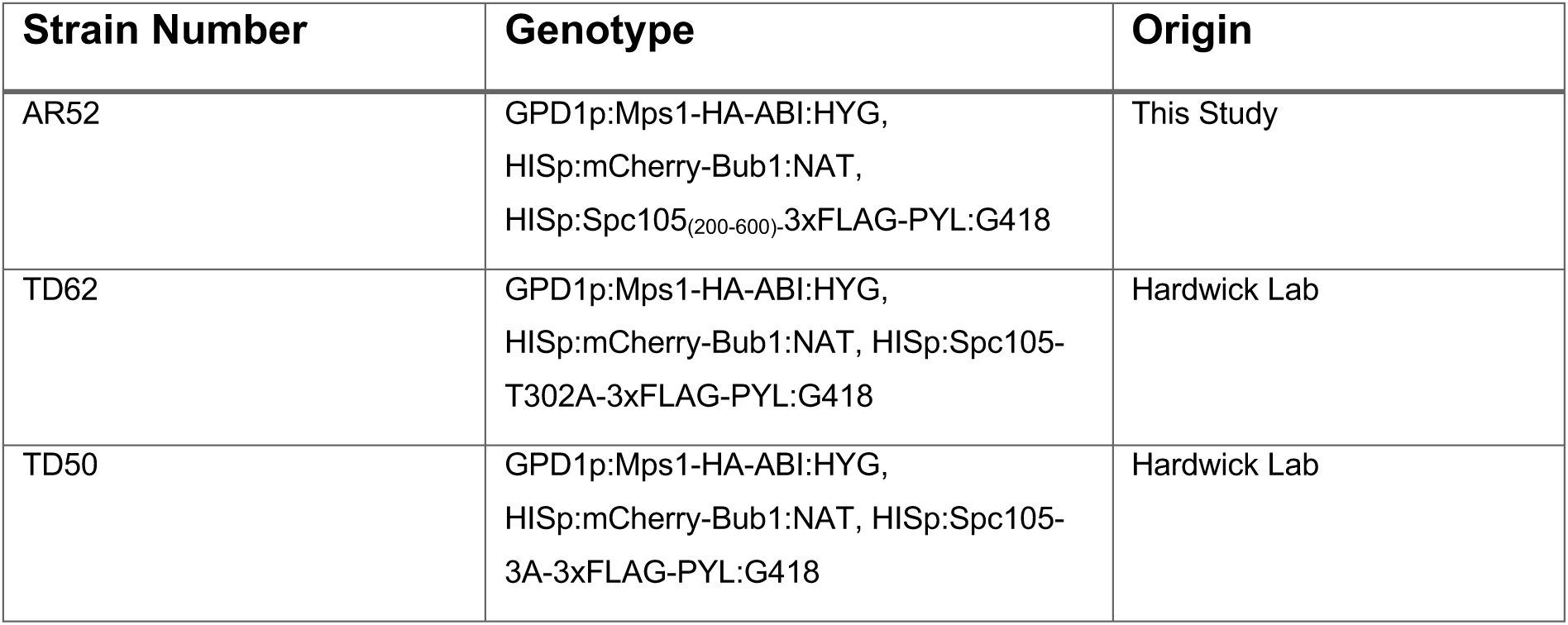
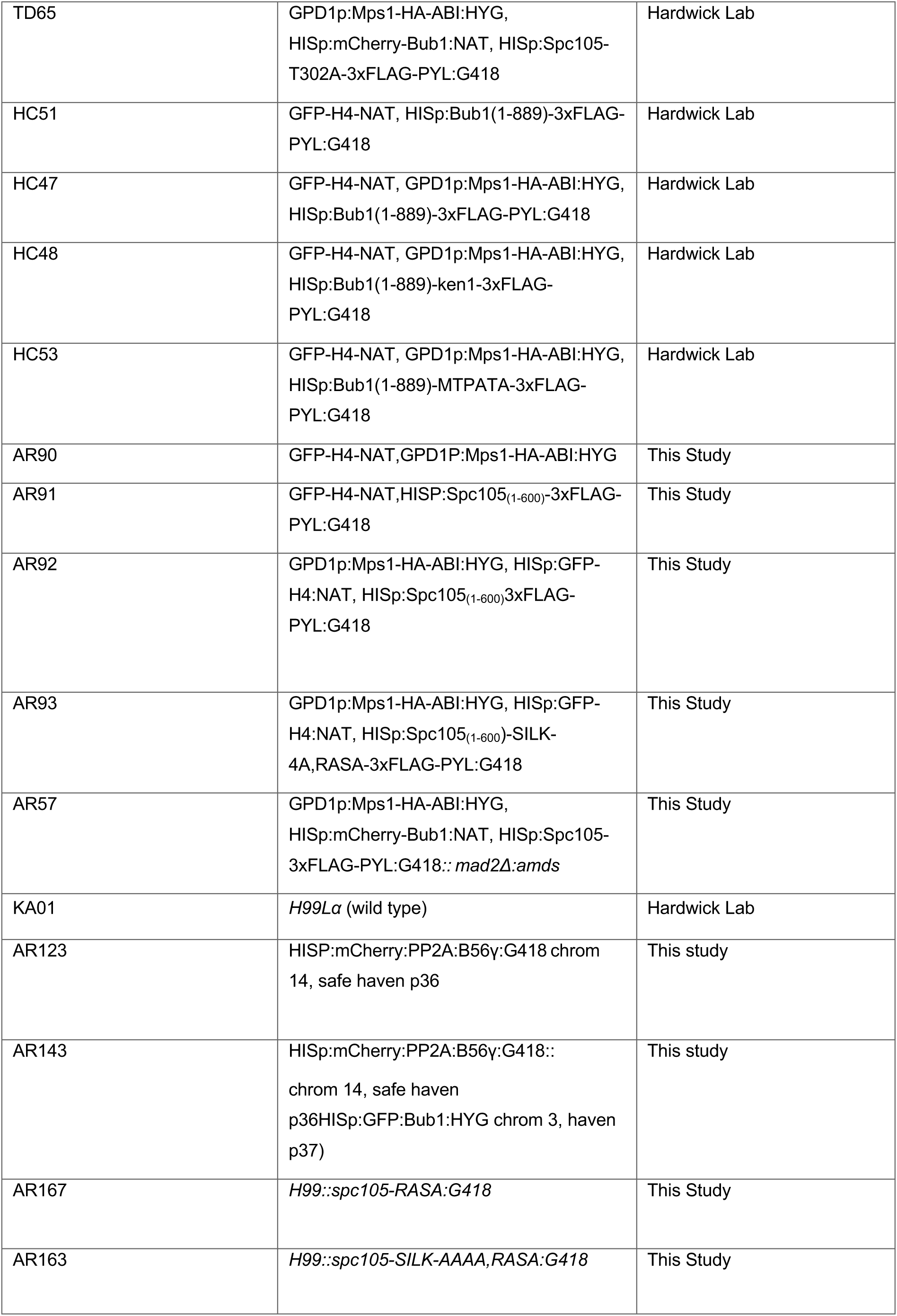
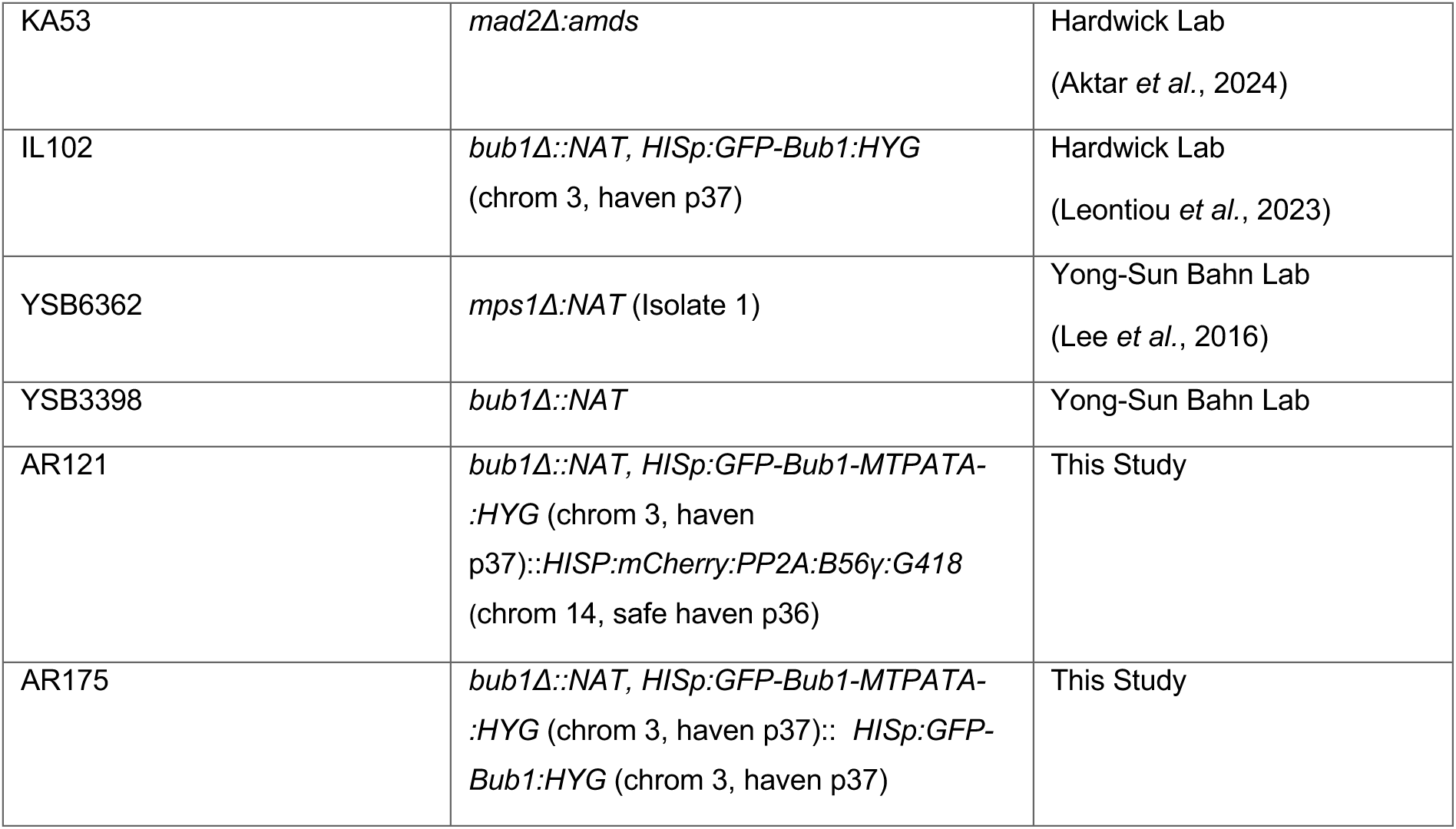

### Plasmid assemblies and site-directed mutagenesis

Generation of point mutations in protein amino acid sequences utilised the QuikChange II Site-Directed Mutagenesis protocol (Aligent). Overlapping primers containing point mutations were designed *via* the QuikChange^®^ Primer Design Web Tool. Initial PCR was carried out according to the manufacturer’s protocol with wild-type plasmid construct templates. Amplified plasmids were incubated with Dpn1 restriction enzyme to degrade methylated wild-type parental DNA, and transformed into competent DH5α *E. coli* cells.

### Construction of ABA-inducible Mps1–ABI and Bub1/Spc105–PYL fusion strains

Fusion constructs for the abscisic acid (ABA)–inducible synthetic checkpoint system were generated by Gibson assembly (NEBuilder HiFi DNA Assembly Master Mix, New England Biolabs) following standard protocols. Truncated coding regions corresponding to the Mps1 kinase domain (residues 474–845), Bub1 (residues 1–889), and Spc105 (residues 1–600) were PCR-amplified from *Cryptococcus neoformans* H99 genomic DNA using Q5 polymerase (New England Biolabs).

PCR fragments were assembled in-frame with the corresponding ABA dimerisation modules and epitope tags to generate Mps1(474–845)–3×HA–ABI, Bub1(1–889)–3×FLAG–PYL, and Spc105(1–600)–3×FLAG–PYL constructs. Each fusion was cloned downstream of either the *GPD1* (CNAG_06699) or Histone H3 (CNAG_06745) constitutive promoter, as appropriate, into plasmids linearised with unique restriction sites to allow directional insertion by Gibson assembly. Assembly reactions were incubated at 50 °C for 1 h, transformed into *E. coli DH5α*, and verified by colony PCR and Sanger sequencing to confirm junction integrity and reading frame.

For mutant constructs, *spc105* alleles carrying SILK→AAAA or RVSF→RASA substitutions and *bub1* carrying the MTPITE→MTPATA mutation were generated by site-directed mutagenesis using overlapping PCR primers and the QuikChange II Site-Directed Mutagenesis protocol (Aligent). The verified mutant fragments were cloned and assembled in the same manner as their wild-type counterparts to produce Spc105(SILK-4A)–PYL, Spc105(RASA)–PYL, and Bub1(MTPATA)–PYL fusion plasmids.

### Primers used in this study

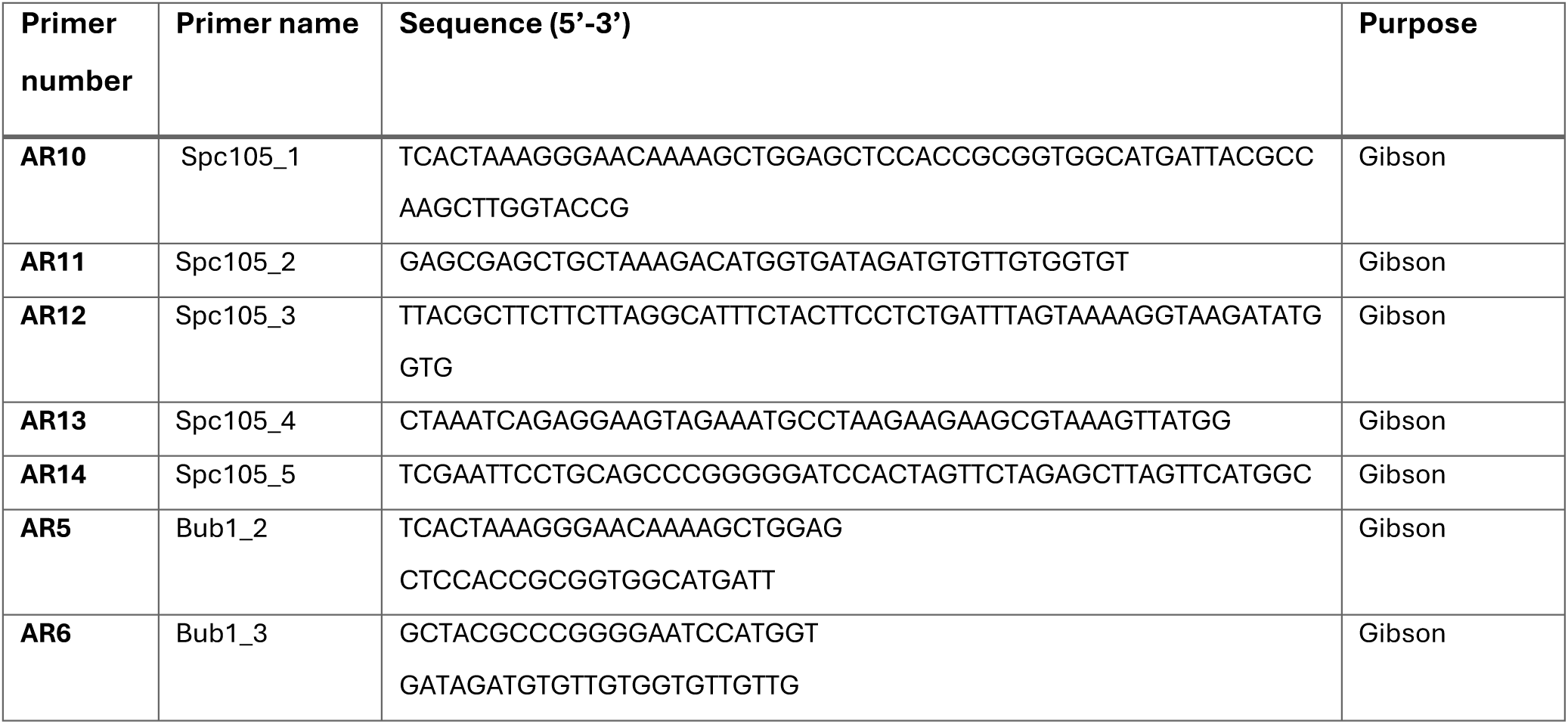
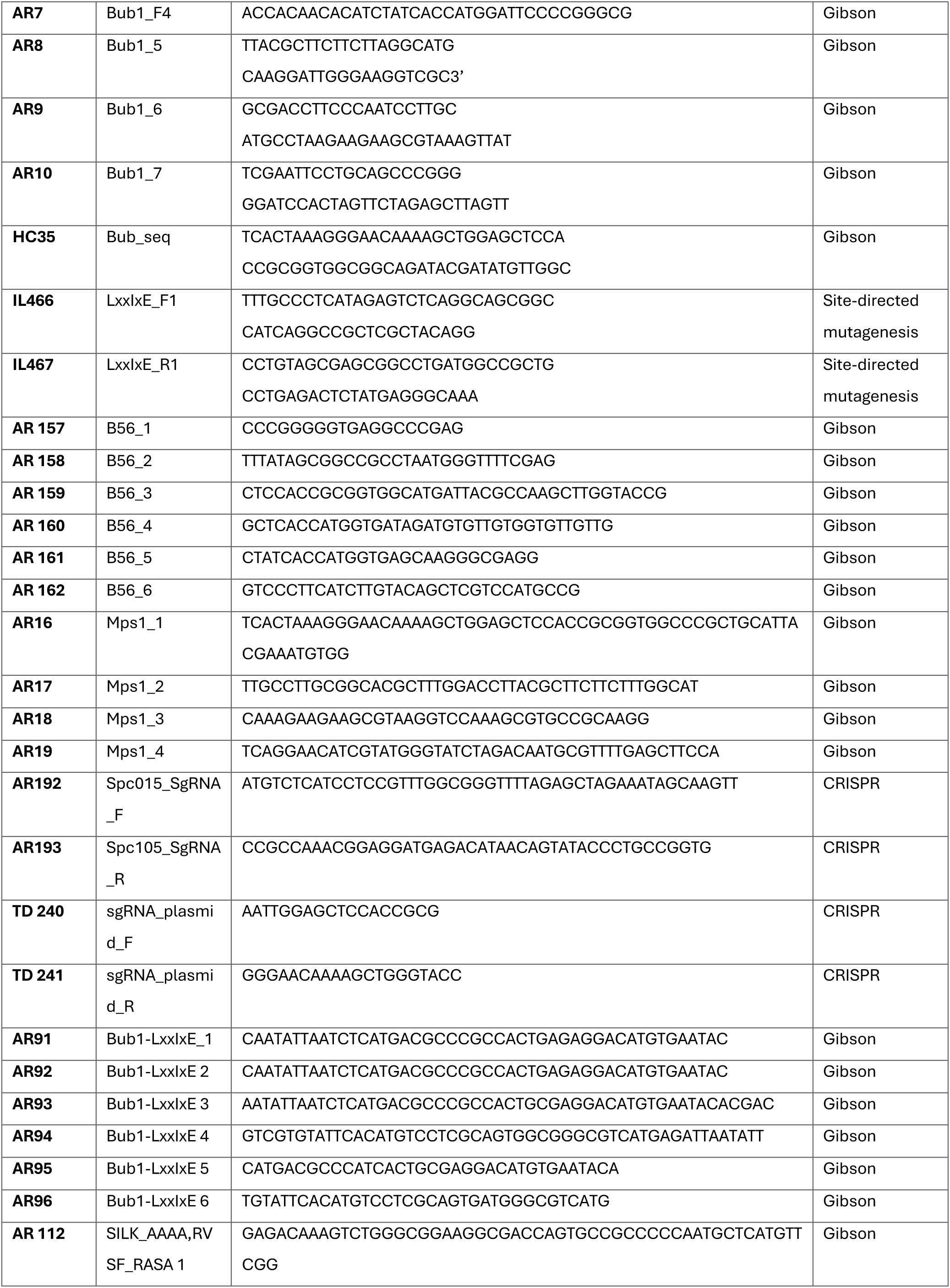
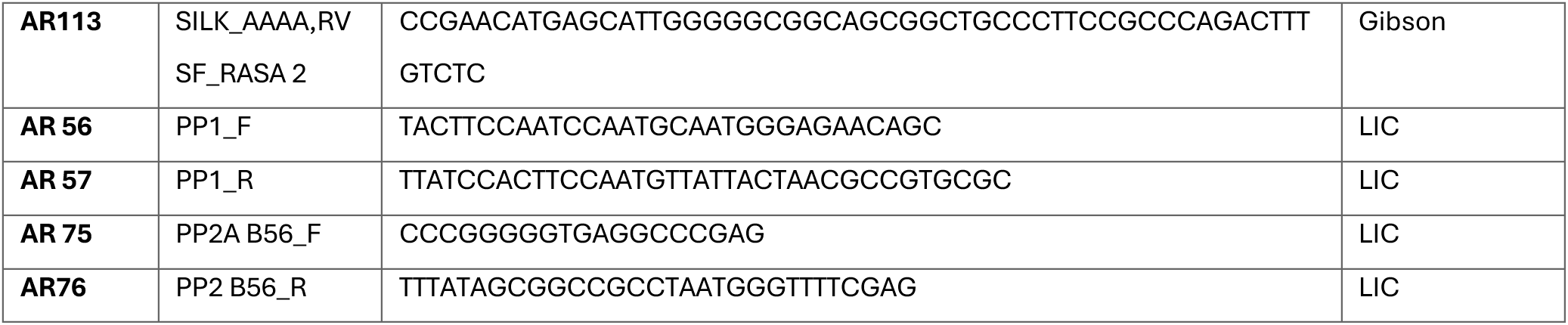

### CRISPR engineering of the *spc105-silk-4A,RASA* strains

Genome editing of *Cryptococcus neoformans* was performed using the CnoCAS9 short-homology CRISPR system previously described (Huang *et al*, 2022) with modifications for precise motif mutagenesis. Briefly, single-guide RNAs (sgRNAs) were designed to target regions flanking the *SPC105* SILK and RVSF motifs. Repair templates were produced by PCR using oligonucleotides carrying alanine-substitution mutations (SILK-AAAA and RVSF-RASA) and ∼50 bp homology arms flanking each site. Cas9–sgRNA and donor DNAs were co-electroporated. Following 3–4 h of recovery in YPDA at 30 °C to allow genomic repair and homologous recombination at the Safe Haven 2 (SH2) locus (Arras et al., 2015), cells were plated on YPDA containing 100 µg mL⁻¹ nourseothricin for selection of transformants. Following 3-4hrs of recovery in YPDA at 30 °C, cells were plated on YPDA + 100 µg mL⁻¹ nourseothricin for selection of homologous recombinants.

Colonies were screened by PCR genotyping of the *spc105* locus, and all edits were verified by Sanger sequencing.

### Synthetic arrest assay with abscisic acid (ABA)

Overnight cultures were diluted to OD600 ∼0.2 and grown for 2 h before treatment with abscisic acid (ABA; 2-cis,4-trans-abscisic acid, Sigma-Aldrich, Cat. 862169) prepared as a 250 mM stock (in DMSO). ABA was added to a final concentration of 250 µM (unless stated otherwise), and cells were incubated for 2 h to induce checkpoint arrest.

### ABA arrest recovery assay

Following ABA-induced arrest, cells were washed five times with 50 ml YES medium and then returned to fresh growth medium for recovery assays as indicated.

### Fluorescence microscopy

Cultures (1–1.5 ml) were centrifuged at 6000 rpm for 1 min, and the resulting pellet was resuspended in 200–500 µL of ice-cold 100% methanol. For imaging, 8 µL of the fixed suspension was applied to a glass slide and covered with a coverslip.

Fixed-cell imaging was performed on a Nikon Ti2 CSU-W1 spinning-disk confocal microscope equipped with a 100× oil immersion Plan Apo VC objective and a Teledyne-Photometrics Prime 95B sCMOS camera. Z-stacks of 11 optical sections were acquired at 0.5 µm intervals using 491 nm (GFP) and 561 nm (mCherry) laser excitation. Exposure times were 300 ms, with laser power minimized to reduce photobleaching. Images were collected with Nikon Elements software.

Image analysis was performed in ImageJ. Brightness and contrast adjustments were carried out in Adobe Photoshop, applied uniformly to entire images and consistently across experimental comparisons.

### CFU viability assays

*C. neoformans* cells were grown to the OD^600^=0.3 in standard conditions, and the cell density at OD^600^ 0.3 was checked with a haemocytometer for calculating the dilutions. Cell cultures of OD^600^ 0.3 were treated with ABA (dissolved in DMSO) at 250 uM or DMSO of the same volume and incubated at 30 °C for 5 h. At relevant timepoints, 0.3 mL of cell cultures were collected *via* centrifuge at 5000 rpm for 3 min. Depending on cell densities at each timepoint as the *C. neoformans* cells duplicate around every 2 h, cultures were serially diluted with ddH_2_O for 10000X −40000X, and 45 µL of cell resuspension containing around 100-200 cells were plated on YPDA agar. The plated cells were incubated at 30 °C for 2 d before colonies were counted, and the cell proliferation rates were calculated as the ratio of initial colony counts, accounting for dilution factors.

### Electroporation transformation

Strains were pre-cultured in YPDA and subsequently diluted into 100 mL fresh medium for overnight growth at 30 °C (25 °C for *bub1*Δ). Cultures were harvested the following morning at an OD^600^ of 0.3–0.36. Cells were collected, washed twice with 50 mL ice-cold water, and once with 50 mL ice-cold electroporation buffer (10 mM Tris-HCl, pH 7.5; 1 mM MgCl₂; 270 mM sucrose). The pellet was resuspended in 35 mL electroporation buffer supplemented with 150 µL of 1 M DTT and incubated on ice for 15 min. Cells were then recovered by centrifugation, washed with 50 mL electroporation buffer, and finally resuspended in ∼100 µL electroporation buffer. For transformation, 40 µI of the cell suspension was gently mixed with ∼4 µg of linearized DNA and transferred to pre-cooled 2 mm electroporation cuvettes. Electroporation was performed using a Bio-Rad Gene Pulser at 1400 V, 600 Ω, and 25 µF. Following the pulse, cells are gently resuspended in YPD and plated onto YPDA plates. The next day, cells were replica-plated onto appropriate selective media. Colonies were screened for correct construct integration by PCR and verified for protein expression by western blotting.

### Serial dilution plating assay

Cells from overnight cultures were diluted to OD^600^ ∼0.4 in distilled water. Ten-fold serial dilutions were spotted onto YPDA plates containing the anti-microtubule drug benomyl (prepared at different concentrations from a 30 mg/ml stock in DMSO) or drug-free controls. Plates were incubated at 30 °C for 48 h. Due to solubility constraints, benomyl stock was added directly to boiling YPD agar.

### Time-lapse microfluidic assays

Microfluidic experiments were performed in 5-chambered Alcatras devices (Crane *et al*, 2014), with chamber heights increased to 7 µm to the *C. neoformans* cell size. Devices were moulded in PDMS from SU8-patterned wafers (Microresist, Berlin). Strain-specific chambers were separated by 2 µm PDMS pillar arrays to prevent intermixing while maintaining identical media conditions. Prior to inoculation, devices were pre-filled with synthetic complete (SC) medium supplemented with 0.2 g/L glucose and 0.05% (w/v) bovine serum albumin (BSA; Sigma-Aldrich) to reduce adhesion. Log-phase cells grown in SC glucose without BSA were then introduced into the chambers. Media flow was maintained at 10 µL/min using an EZ-Flow system (Fluigent). Cells were cultured in SC for 5 h before switching to SC medium containing 2.5 µg/mL nocodazole. After 3 h of nocodazole treatment, the drug was removed (8 h total elapsed), and cells were returned to fresh SC medium for a further 12 h. Cy5 dye (Abcam) was included in the media containing nocodazole to monitor media switching events. Imaging continued throughout, giving a total experimental duration of 20 h. Image acquisition was carried out at two stage positions per strain using a Nikon Ti-E epifluorescence microscope equipped with a 60× oil immersion objective (NA 1.4), a Prime95B sCMOS camera (Teledyne Photometrics), and OptoLED illumination (Cairn Research). Z-stacks of five sections at 0.6 µm spacing were collected at 2 minute intervals..

### Time-lapse image processing and analysis

Cell segmentation was performed using a convolutional neural network developed by the Swain laboratory (Clark *et al*, 2025; Pietsch *et al*, 2023). Further analysis and visualisation were carried out using custom Python software (code is available on request). For quantification of GFP-Bub1 kinetochore dwell time, maximum-intensity projections were generated from GFP Z-stacks. The ratio of the median fluorescence intensity of the five brightest pixels within each segmented cell to the median whole-cell intensity was calculated, providing a robust measure of Bub1 accumulation at kinetochores. Outlier cells (with values >3 scaled median absolute deviations from the median) and cells missing for part of the time course were excluded. Ratios were normalised to the mean of all wild-type values in the absence of nocodazole. A threshold of 1.1 was used to define Bub1 kinetochore localisation at each time point. Brief signal losses due to focus shifts were corrected using 1D morphological closing of thresholded data with a three-pixel structuring element. The mean lengths of localization above the threshold are presented for the period from 11 h 40 minutes until the end of the acquisition. For heat map visualisations, 40 cells per strain were randomly selected from those tracked for the entire experiment.

### Immunoblot analysis

For protein extracts, 10 mL cultures (OD^600^ ∼0.5) were harvested by centrifugation, washed once, and snap-frozen in liquid nitrogen. Pellets were resuspended in 2× SDS sample buffer containing 200 mM DTT and 1 mM PMSF. Cells were disrupted with 0.5 mm Zirconia/Silica beads (Thistle Scientific) in a multi-bead beater (BioSpec Products) for 1 min. Lysates were clarified by centrifugation, boiled for 5 min at 95 °C, and centrifuged again for 5 min at 13,000 rpm. Cleared extracts were separated by SDS–PAGE and transferred to nitrocellulose membranes (Amersham Protran 0.2 µm, GE Healthcare Lifescience) using a semi-dry transfer system (TE77, Hoefer) in buffer containing 25 mM Tris, 130 mM glycine, and 20% methanol. Transfers were run for 1.5–2 h at 150–220 mA. Ponceau S staining was used to verify transfer efficiency. Membranes were blocked in PBS + 0.04% Tween-20 with 4% Marvel skimmed milk powder for ≥30 min at room temperature, then incubated overnight at 4 °C with primary antibodies (anti-FLAG, anti-mCherry, anti-GFP, or anti-HA; all 1:1000) diluted in blocking buffer. After three 10 min washes in PBS + 0.04% Tween-20, membranes were incubated with HRP-conjugated secondary antibodies (1:5000) for 1 h at room temperature, washed again, and developed by chemiluminescence (SuperSignal West Pico or SuperSignal West Femto, Thermo Fisher Scientific).

### Lysis of large-scale cell extracts and co-immunoprecipitation

*Cryptococcus* cells were cultured in 500 mL YPDA to an OD600 of ∼0.5. Cultures were treated with 2.5 µg/mL nocodazole for 3 h to induce spindle checkpoint arrest. Cells were harvested by centrifugation at 5000 rpm for 15 min at 4 °C, and the resulting pellets were frozen dropwise in liquid nitrogen. Frozen pellets were ground manually to a fine powder using a pre-chilled mortar and pestle. Cell powders were resuspended in lysis buffer (50 mM HEPES pH 7.6, 75 mM KCl, 1 mM MgCl₂, 1 mM EGTA, 10% glycerol, 0.1% Triton X-100, 1 mM Na₃VO₄, 10 µg/ml CLAAPE protease inhibitor cocktail [chymostatin, leupeptin, aprotinin, antipain, pepstatin, E-64; dissolved in DMSO at 10 mg/mL each], 1 mM PMSF, 0.01 mM microcystin). One gram of yeast powder was resuspended in 1 mL lysis buffer. Samples were sonicated on ice using 5 s on / 5 s off cycles for 1 min. Lysates were clarified by centrifugation at 22,000 rpm for 30 min at 4 °C, and the supernatant was incubated with GFP-TRAP magnetic agarose beads (ChromoTek) for 1 h at 4 °C. Beads were washed at least nine times in wash buffer (50 mM HEPES pH 7.6, 75 mM KCl, 1 mM MgCl₂, 1 mM EGTA, 10% glycerol) followed by one wash in PBS containing 0.001% Tween-20. Bound proteins were eluted by boiling the beads in 2× SDS sample buffer supplemented with 200 mM DTT for 5–10 min at 95 °C, and analysed by SDS–PAGE.

### Recombinant protein purification

The C-terminal kinase domain of *C. neoformans* Mps1 (residues 478–784) was expressed as an N-terminal His₆–MBP fusion in *E. coli* BL21 (DE3) cells. Cultures were grown at 37 °C to an OD^600^ of ∼0.6 and induced with 0.3 mM IPTG at 18 °C overnight. Cells were harvested by centrifugation (5,000 × g, 10 min, 4 °C) and resuspended in lysis buffer containing 20 mM Tris–HCl (pH 8.0), 125 mM NaCl, 4 mM imidazole, and 1 mM β-mercaptoethanol (β-ME). Lysis was performed by sonication on ice, and clarified lysates were obtained by centrifugation at 30,000 × g for 30 min at 4 °C. The supernatant was applied to Ni²⁺–NTA agarose resin (Qiagen) pre-equilibrated in lysis buffer. After extensive washing, bound proteins were eluted using 20 mM Tris–HCl (pH 8.0), 125 mM NaCl, 200 mM imidazole, and 1 mM β-ME. The eluted fractions containing His–MBP–Mps1 were pooled and dialysed overnight at 4 °C against 20 mM Tris–HCl (pH 7.0), 125 mM NaCl, 5% (v/v) glycerol, and 2 mM DTT. To remove contaminants and exchange buffer, the dialysed sample was further purified by ion-exchange chromatography (IEX) using a Mono Q column (Cytiva) equilibrated in 20 mM Tris–HCl (pH 7.0), 1 M NaCl, 5% glycerol, 2 mM DTT, followed by size-exclusion chromatography (SEC) on a Superdex 200 Increase 10/300 GL column (Cytiva) equilibrated in 20 mM Tris–HCl (pH 7.0), 125 mM NaCl, and 2 mM DTT. Peak fractions corresponding to monomeric His–MBP–Mps1 were pooled, concentrated, flash-frozen in liquid nitrogen, and stored at –80 °C until use in kinase assay.

The N-terminal region of *C. neoformans* Spc105 (residues 1–600) was expressed as an N-terminal His_6_–MBP fusion in E. coli BL21 (DE3) cells. Cultures were grown at 37 °C to an OD^600^ of ∼0.6 and induced with 0.3 mM IPTG at 18 °C overnight. Cells were harvested by centrifugation (5,000 × g, 10 min, 4 °C) and resuspended in lysis buffer containing 50 mM Tris–HCl (pH 7.0), 500 mM NaCl, 5 mM imidazole, and 1 mM β-mercaptoethanol (β-ME). Lysis was performed by sonication on ice, and lysates were clarified by centrifugation at 30,000 × g for 30 min at 4 °C. The supernatant was incubated with Ni²⁺–NTA agarose resin (Qiagen) pre-equilibrated in lysis buffer. After extensive washing, bound proteins were eluted with 50 mM Tris–HCl (pH 7.0), 500 mM NaCl, 250 mM imidazole, and 1 mM β-ME. Eluted fractions containing His–MBP–Spc105 were pooled and dialysed overnight at 4 °C against 50 mM Tris–HCl (pH 8.0), 250 mM NaCl, 5% (v/v) glycerol, and 2 mM DTT.

### Phospho-mass spectrometry

*In vitro* phosphorylation assays were performed using purified *C. neoformans* Mps1 kinase and recombinant Spc105 substrate. Reactions contained 1.5 µg Mps1 and 6 µg Spc105 in 20 µL kinase buffer comprising 20 mM HEPES (pH 7.6), 100 mM KCl, 10 mM MgCl₂, 1 mM DTT, and 100 µM ATP (Thermo Fisher Scientific). Proteins from kinase reactions were resolved by SDS–PAGE (NuPAGE Novex 4–12% Bis-Tris gel, Life Technologies) using MES buffer and visualised with InstantBlue™ stain (Abcam). Gel bands corresponding to Spc105 were excised, de-stained in 50 mM ammonium bicarbonate and 100% (v/v) acetonitrile, reduced with 10 mM dithiothreitol (30 min, 37 °C), and alkylated with 55 mM iodoacetamide (20 min, room temperature, dark). Samples were digested overnight at 37 °C with trypsin (12.5 ng µL⁻¹), diluted 1:1 with 0.1% TFA, and desalted using StageTips. Peptides were eluted in 40 µL 80% acetonitrile/0.1% TFA, concentrated by vacuum centrifugation, and resuspended in 0.1% TFA prior to LC–MS/MS analysis.

Peptides were analysed on an Orbitrap Fusion™ Lumos™ Tribrid™ mass spectrometer (Thermo Fisher Scientific) coupled to an Ultimate 3000 HPLC (Dionex). Separation was achieved on a 50 cm EASY-Spray column (2 µm, Thermo Scientific) operated at 50 °C using a gradient of 2–40% solvent B (80% acetonitrile, 0.1% formic acid) over 150 min, followed by a ramp to 95% B in 11 min at 0.25 µL min⁻¹. Survey scans were collected at 120,000 resolution (m/z 350–1500) with an ion target of 4.0 × 10⁵ and injection time of 50 ms. MS² spectra were acquired in the ion trap using HCD fragmentation (NCE = 28), an isolation window of 1.4 Th, ion target of 2.0 × 10⁴, and 60 s dynamic exclusion.

Raw data were processed with MaxQuant (v1.6.1.0) using the Andromeda search engine against an in-house *C. neoformans* var. *grubii* H99 protein database. Peptide tolerances were 20 ppm (first search) and 4.5 ppm (main search). Carbamidomethylation (Cys) was set as a fixed modification, and oxidation (Met) and phosphorylation (Ser/Thr/Tyr) as variable modifications. Trypsin specificity was enforced with up to two missed cleavages permitted. Label-free quantification employed the MaxLFQ algorithm, and peptide/protein identifications were filtered to 1% FDR. Statistical analyses were performed using Perseus (v1.6.2.1).

### γ-³²P-ATP kinase assays

For radiolabelled assays, ∼500 kBq γ-³²P-ATP (PerkinElmer) was added to kinase reactions, as detailed above, prior to incubation. After electrophoresis, gels were Coomassie-stained, dried on a Hoefer™ slab gel dryer, and exposed to autoradiography film or a BAS phosphorimaging screen for quantification on a Typhoon™ biomolecular imager.

